# Facilitating candidate gene discovery in an emerging model plant lineage: Transcriptomic and genomic resources for *Thalictrum* (Ranunculaceae)

**DOI:** 10.1101/2020.06.25.171215

**Authors:** Tatiana Arias, Diego Mauricio Riaño-Pachón, Verónica S. Di Stilio

## Abstract

The plant genus *Thalictrum* is a representative of the order Ranunculales (a sister lineage to all other Eudicots) with diverse floral morphologies, encompassing four sexual systems and two pollination modes. Previous studies suggest multiple transitions from insect to wind pollination within this genus, in association with polyploidy and unisexual flowers, but the underlying genes remain unknown. We generated a draft reference genome for *Thalictrum thalictroides*, a representative of a clade with ancestral floral traits (diploidy, hermaphroditism, and insect pollination) and a model for functional studies. To facilitate candidate gene discovery in flowers with different sexual and pollination systems we also generated floral transcriptomes of *T. thalictroides* and of wind-pollinated, andromonoecious (staminate and hermaphroditic flowers on the same plant) *T. hernandezii*.

The *T. thalictroides* draft genome assembly consisted of 44,860 contigs (N50=12,761 bp. and 243 Mbp. total length) and contained 84.5% conserved embryophyte single-copy genes. Floral transcriptomes from Illumina sequencing and *de novo* assembly contained representatives of most eukaryotic core genes (approximately 80%), with most of their genes falling into common orthologous groups (orthogroups). Simple Sequence Repeat (SSR) motifs were also identified, which together with the single-copy genes constitute a resource for population-level or phylogenetic studies. Finally, to validate the utility of these resources, putative candidate genes were identified for the different floral morphologies using stepwise dataset comparisons. In conclusion, we present genomic and transcriptomic resources for *Thalictrum*, including the first genome of *T. thalictroides* and potential candidate genes for flowers with distinct sexual and pollination systems.

## INTRODUCTION

Genomic resources are a valuable tool for studying development within an evolutionary context, as they enable the search for candidate loci and their regulatory regions, contributing to the upgrading of study systems into model lineages (Chang and Cook, 2002; Di Stilio et al., 2017). RNA-Seq enables the study of most genes expressed in a sample and the comparison of gene expression levels across developmental stages, conditions, populations or species (Wolf, 2013). The success of this technique to address evolutionary questions relies on its applicability to nonestablished model organisms and its growing toolkit and bioinformatics support (Liu et al., 2015).

The plant genus *Thalictrum* is an emerging model lineage within the Ranunculales (Damerval and Becker, 2017), the sister group to all other eudicots (a monophyletic group of dicotyledonous plants) (Lane et al., 2018). The genus is therefore an extant representative of a privileged phylogenetic node, before a major evolutionary radiation within the flowering plants (Zeng et al., 2017). *Thalictrum* is a temperate genus of circa 200 species with floral diversity that encompasses unisexual and wind-pollinated flowers, in association with polyploidy (Soza et al., 2012, 2013). Certain species, mainly *T. thalictroides*, are amenable to virus-induced gene silencing (Di Stilio et al., 2010), which has enabled functional studies of ABC model floral MADS-box genes (Di Stilio et al., 2013; Galimba and Di Stilio, 2015; Galimba et al., 2012, 2018; Soza et al., 2016). Two chloroplast genomes have been assembled to date for the genus, for *T. coreanum* (Park et al., 2015) and *T. thalictroides* (from this dataset, Morales Briones et al., 2019).

Here, we contribute genomic resources for *Thalictrum*, including the first *de novo* draft genome of *Thalictrum thalictroides*, a diploid, hermaphroditic and insect-pollinated species, and two *de novo* floral transcriptomes, one for *T. thalictroides* and the other for tetraploid, andromonoecious and wind-pollinated *T. hernandezii*. A qualitative comparison amongst these transcriptomes provides a preliminary list of enriched transcription factor families and potential candidate genes for their distinct phenotypes. Microsatellite markers were also identified as a resource for population-level genetic studies. These genetic resources represent a tool to facilitate the elucidation of the genetic underpinnings of floral diversity in *Thalictrum*, and across the order Ranunculales.

## METHODS

### 1. *Thalictrum thalictroides* nuclear genome

#### Plant materials

Two live accessions of *Thalictrum thalictroides* (Tt), TtWT478 and TtWT964, were grown in the University of Washington greenhouse (Suppl. Table 1).

#### DNA extraction

Total DNA was extracted from healthy young leaves of the two individuals using a standard CTAB method with modifications (Doyle and Doyle, 1987; Shyu and Hu, 2013). DNA quality and concentration were measured in a Qubit^®^ (Life Technologies, Carlsbad, CA, USA) and with agarose gel electrophoresis. Short read sequencing was carried out by Beijing Genomics Institute (BGI, Hong Kong).

#### DNA Library preparation

Total genomic DNA was sheared by sonication for 15–24 min using a Bioruptor (Diagenode, Inc., New Jersey, USA). Size-selected samples (200–400 bp) were extracted using x-tracta™ Gel Extraction tools (USA Scientific, Ocala, Florida, USA) and purified with Gel DNA Purification Kit (Qiagen, Germantown, Maryland, USA) for the endrepair, adenylation of 3’ ends, ligation, and enrichment steps. Sheared DNA was visualized on a 2% low-melting-point agarose gel stained with ethidium bromide. Illumina’s TruSeq paired-end (2 x 100 bp) libraries were generated for each individual with three different insert sizes (170 bp, 500 bp and 800 bp), following manufacturer’s instructions. Libraries were assessed in an Agilent 2100 bioanalyzer with the DNA 1000 Kit (Agilent, Santa Clara, California) to determine average length, quantitated by real-time quantitative PCR (qPCR) using a TaqMan^®^ Probe and amplified off-board on a cBot (TruSeq PE Cluster Kit v3–cBot–HS, Illumina) to generate clusters on the sequencing flow cell, then sequenced on a HiSeq 2000 (TruSeq SBS Kit v3-HS, Illumina).

#### Read pre-processing, genome assembly and annotation

Raw read quality was visually inspected with FastQC (Babraham Bioinformatics) and then treated with Trimmomatic v0.38 (Bolger et al., 2014) to remove adapter sequences and low-quality bases, keeping only paired-end reads with at least 80 bp for *de novo* assembly. Initial *de novo* genome assemblies were conducted in CLC Genomics Workbench v7.5.1, SOAP *de novo* v2.04 (Luo et al., 2012b) and MaSuRCA v3.2 (Zimin et al., 2013). Assemblies were scanned to identify contigs originating from contaminants using BlobTools (Laetsch and Blaxter, 2017). The genome sequences of potential contaminants were downloaded from NCBI and used to map the clean reads, using BBDuk (Bushnell, 2017), keeping only unmapped reads. Final assemblies were generated with MaSuRCA v3.2.9 (Zimin et al., 2013). One assembly was generated for each of the two *T. thalictroides* accessions, and a third was generated by combining the data from the two accessions. Each assembly was polished using Pilon (Walker et al., 2014), with two rounds of error correction. Assembly statistics were computed with Quast v5.0.2 (Gurevich et al., 2013) and sequence repeats were identified with Repeat Modeler v1.0.11 (Smit and Hubley, 2008) and Repeat Masker v4-0-9-p2 (Smit et al., 2013). RNASeq datasets generated from *T. thalictroides* flowers (see below), together with a publicly-available 1KP project transcriptome (Matasci et al., 2014) (http://www.onekp.com), and predicted protein sequences from the *Aquilegia coerulea* genome (Filiault et al., 2018) were used as extrinsic evidence for protein-coding gene prediction using Braker2 (Hoff et al.). RNASeq data were also employed for protein-coding gene prediction with StringTie (Pertea et al., 2015) and Transdecoder (Haas et al., 2013); *ab initio* protein-coding gene prediction was generated with SNAP (Korf, 2004) using *Arabidopsis thaliana* as a model. Final models for protein-coding genes were produced with EVidenceModeler (Haas et al., 2008) with the following weights: *Ab initio* prediction (SNAP): 1; Predictions based on RNA-Seq and Protein alignments (Braker2): 10; Predictions based on RNASeq alone (StringTie): 5; Protein alignments alone (GenomeThreader): 3 and Genes identified/predicted by BUSCO: 10. tRNAs were predicted with tRNAscan-SE v2.0 (Lowe and Eddy) and ribosomal RNA genes with RNAmmer v1.2 (Lagesen et al., 2007). Other non-protein-coding RNA (ncRNA) were predicted with Infernal v1.1.2 (Nawrocki and Eddy, 2013) and Rfam v14.1 (Kalvari et al., 2018). Predicted proteincoding genes were evaluated with TransRate (Smith-Unna et al., 2016) against *Papaver somniferum* (Guo et al., 2018) and *Aquilegia coerulea* (Filiault et al., 2018) (Ranunculales); *Arabidopsis thaliana* (Cheng et al., 2017) and *Solanum lycopersicum* (The Tomato Genome Consortium et al., 2012), representing the two main eudicot orders. Functional annotation was conducted with BLASTp searches against the *Arabidopsis thaliana* proteome and the SwissProt and TrEMBL protein databases. Protein functional descriptions and gene ontology terms were added with the “Automated Assignment of Human Readable Descriptions” (AHRD) tool (The Tomato Genome Consortium et al., 2012). Conserved regions, domains, and sequence features were identified on protein sequences with InterProScan v5.35-74.0 (Jones et al., 2014; Mitchell et al., 2019). Transcription-associated proteins (TAPs) were identified using the approach described for the construction of PlnTFDB v3.0 (http://plntfdb.bio.uni-potsdam.de) (Pérez-Rodríguez et al., 2010; Riaño-Pachón et al., 2007). Classification rules and software to assign protein sequences into TAP families based on their domain architecture was obtained from https://bitbucket.org/diriano/mytfdb.

Benchmarking Universal Single-Copy Orthologs (BUSCO) v3.0 (Waterhouse et al., 2018) were used to quantify the completeness of the genome assemblies, annotated gene sets, and transcriptomes based on evolutionarily-informed expectations of gene content from near-universal single-copy orthologs selected from OrthoDB v9.0 (Lopez et al., 2017). Clusters of orthologous genes (orthogroups) were identified with OrthoFinder v2.3.3 (Emms and Kelly, 2015). Sequence similarity was computed with Diamond v0.9.24 (Buchfink et al., 2014) and gene clusters were generated with MCL v14-137 (Dongen, S., 2000), with 1.5 inflation.

### 2. RNA-Seq of *Thalictrum thalictroides* and *T. hernandezii* floral transcriptomes

#### Plant materials

Fresh open flowers at the same developmental stage were collected from an individual of *T. thalictroides* that was also used for genome sequencing (TtWT478, hermaphroditic flowers) and from *T. hernandezii* (Th_HWT441 hermaphroditic and Th_SWT441 staminate flowers) (Suppl. Table 1).

#### RNA extractions

Flowers of *T. thalictroides* and *T. hernandezii* were flash-frozen in liquid nitrogen and total RNA was extracted with Trizol (Invitrogen, Carlsbad, California, USA), following manufacturer’s instruction. RNA quality (RIN ≥ 6.5) and concentration (≥ 250 ng/μL) was determined in an Agilent 2100 bioanalyzer (Agilent Technologies, Palo Alto, California, USA) combined with agarose gel electrophoresis and a NanoDrop (Thermo Scientific, Waltham, Massachusetts, USA).

#### RNA Library preparation

Three flower libraries were built: *T. thalictroides* hermaphrodite (Tt_H), *T. hernandezii* hermaphrodite (Th_H) and *T. hernandezii* staminate (Th_S). A mixed floral stage library of *T. thalictroides* (1KP, GBVZ, Matasci et al., 2014) was also included in the analysis. RNA-seq was carried out by Beijing Genomics Institute (BGI, Hong Kong).

#### Read pre-processing and Sequence assembly

The data was cleaned to eliminate potential contaminants, assembled *de novo* in Trinity v2.8.5 (Grabherr et al., 2011) and assessed for completeness with BUSCO v3.0 (as for the genome assembly). Only transcripts assigned to Viridiplantae (green plants) were used in further analyses. Two *de novo* transcriptome assemblies were generated: one for *T. thalictroides* and another for *T. hernandezii*, joining the RNASeq libraries for the two types of flowers in the latter. Only contigs larger than 200 bp were used in further analyses. Assembled transcripts were compared against NCBI’s nr protein database using Diamond v0.9.23 and assessed in MEGAN-LR v6.14.2.

#### Annotations

Polypeptides encoded by the assembled transcripts were identified with TransDecoder v5.5.0 (https://transdecoder.github.io/) using default parameters, including BLASTp hits against SwissProt and HMM hits against PFAM. Functional annotation, including identification of TAPs and clustering of shared (orthologous) genes, or “orthogroups”, was carried out as for the genome annotation.

#### Identification of genes involved in insect vs wind pollination, and male vs hermaphrodite flower morphologies

First, we performed a three-way comparison to identify orthogroups amongst the *T. thalictroides* genome and the *T. thalictroides* and *T. hernandezii* de novo transcriptomes. We considered that orthogroups present in the *T. thalictroides* genome and transcriptome, but not found in the *T. hernandezii* transcriptome, are more likely to contain high-confidence genes involved in the development of floral traits that characterize insect-pollinated flowers. Conversely, orthogroups expressed exclusively in *T. hernandezii* (not found in the *T. thalictroides* transcriptome) that can similarly be mapped onto the *T. thalictroides* reference genome, were considered more likely to contain high-confidence genes involved in the development of floral traits that characterize wind-pollinated flowers.

Second, we further analyzed the *T. hernandezii*-specific set of orthogroups described above to identify transcripts associated with the different floral sexes present within *T. hernandezii*, staminate (male) vs. hermaphrodite. To that end, we computed the expression level of the *de novo* assembled *T. hernandezii* transcripts in the two *T. hernandezii* datasets (Ther_S and Ther_H) using Salmon v0.14.0 (Patro et al., 2017) with options “--seqBias --validateMappings --recoverOrphans --libType A” and considered a transcript as expressed when it had at least a single mapped read. For the data intersections of interest, we performed an enrichment analysis of the families of TAPs using a Fisher’s Exact Test, correcting p-values for False Discovery Rate (FDR) with the BH method (Benjamini and Hochberg, 1995). Similarly, we performed Gene Ontology (GO) term enrichment analysis, using topGO (Alexa and Rahnenfuhrer, 2010), with the weight0 method, correcting p-values for FDR using BH.

#### SSR markers search

Simple sequence repeat (SSR) markers and loci with di- to hexa-nucleotide repeats were identified in the genome and transcriptomes using the MicroSAtellite Identification tool (MISA) (Thiel et al., 2003).

#### Data availability

Raw reads and assemblies for the genome and transcriptomes from this study have been deposited in NCBI as BioProject PRJNA439007 and Sequence Read Archive SRP136081 (Suppl. Table 2). Additional data can be found in figshare (https://figshare.com):

Go enrichment: https://doi.org/10.6084/m9.figshare.12465254.v1
MISA: https://doi.org/10.6084/m9.figshare.11984370.v8
Orthogroups: https://doi.org/10.6084/m9.figshare.11984358.v3
TAPs: https://doi.org/10.6084/m9.figshare.11984313.v1

## RESULTS AND DISCUSSION

Large-scale genomics data were generated for *T. thalictroides*, and transcriptomes for *T. thalictroides* and *T. hernandezii* (Ranunculaceae), two species with distinct floral morphologies (Fig. 1). The results presented here bring the known set of protein-coding genes for this genus to 33,624 (predicted from genome sequence, Bioproject: PRJNA439007), from approximately 132 nuclear genes, 10,461 ESTs and 130 population sets (NCBI).

**FIGURE 1.**
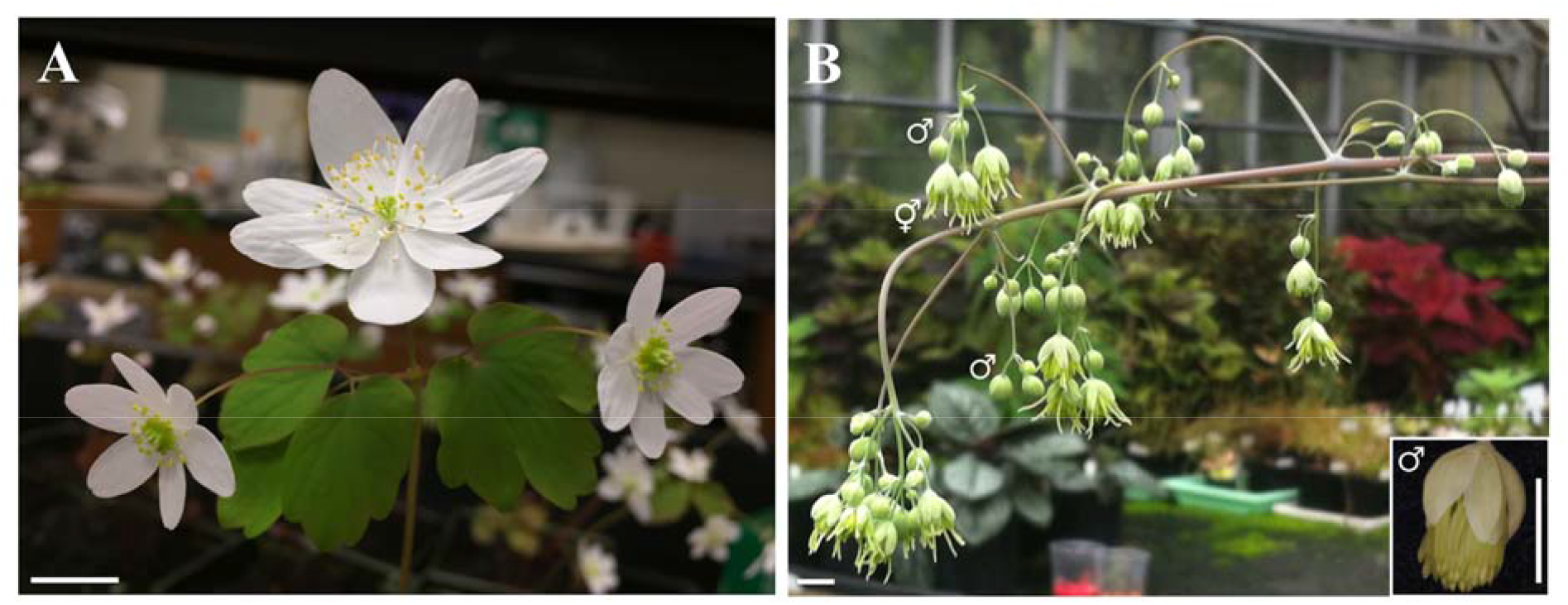
Floral phenotypes of *Thalictrum* study species. Inflorescence of (A) Hermaphroditic *T. thalictroides* with hermaphrodite open flowers and (B) Andromonoecious *T. hernandezii* with staminate buds (♂) and hermaphrodite open flowers (□) occurring together in an inflorescence; inset, detail of young staminate flower. Scale bar = 1 cm.

### A draft nuclear genome for *Thalictrum thalictroides*

#### Genome sequencing and de novo assembly

Paired-end sequencing of six libraries from two live accessions of *T. thalictroides* (TtWT478 and TtWT964) resulted in 49,105,897 sequenced fragments, after quality-trimming and contaminant removal (Suppl. Table 2). On average, 11.6 million reads were generated per library. The *T. thalictroides* draft reference genome assembly consisted of 44,860 contigs with N50=12,761 bp (Table 1; Suppl. Table 3) and contained 83.8% conserved embryophyte single-copy genes (Fig. 2), this estimate increased to 84.5% after gene prediction. These genome completeness results are within-range of other draft genomes of non-traditional species, such as *Calotropis gigantea* (77-88%, Hoopes et al., 2017) and certain Brassicaceae (61-95%, Lopez et al., 2017), yet lower than the reference genomes used in our orthogroup analysis (94.5%-99.6%, Suppl Table 8).

**TABLE 1.** Summary statistics for *de novo* genome assemblies of *Thalictrum thalictroides*. WT964 and WT 478 are the two sequenced accessions.

|  | Metric | Consensus | WT964 | WT478 |
| --- | --- | --- | --- | --- |
| Assembly statistics<br>(Quast v5.0.2) | Number of contigs | 44,860 | 41,892 | 41,384 |
|  | Total length (bp) | 243,091,176 | 205,112,749 | 167,308,766 |
|  | Largest contig (bp) | 119,890 | 80,531 | 57,549 |
|  | GC (%) | 36.23 | 36.29 | 36.61 |
|  | N50 (bp) | 12,761 | 10,189 | 8,017 |
|  | L50 | 4,281 | 4,739 | 4,875 |

**FIGURE 2.**
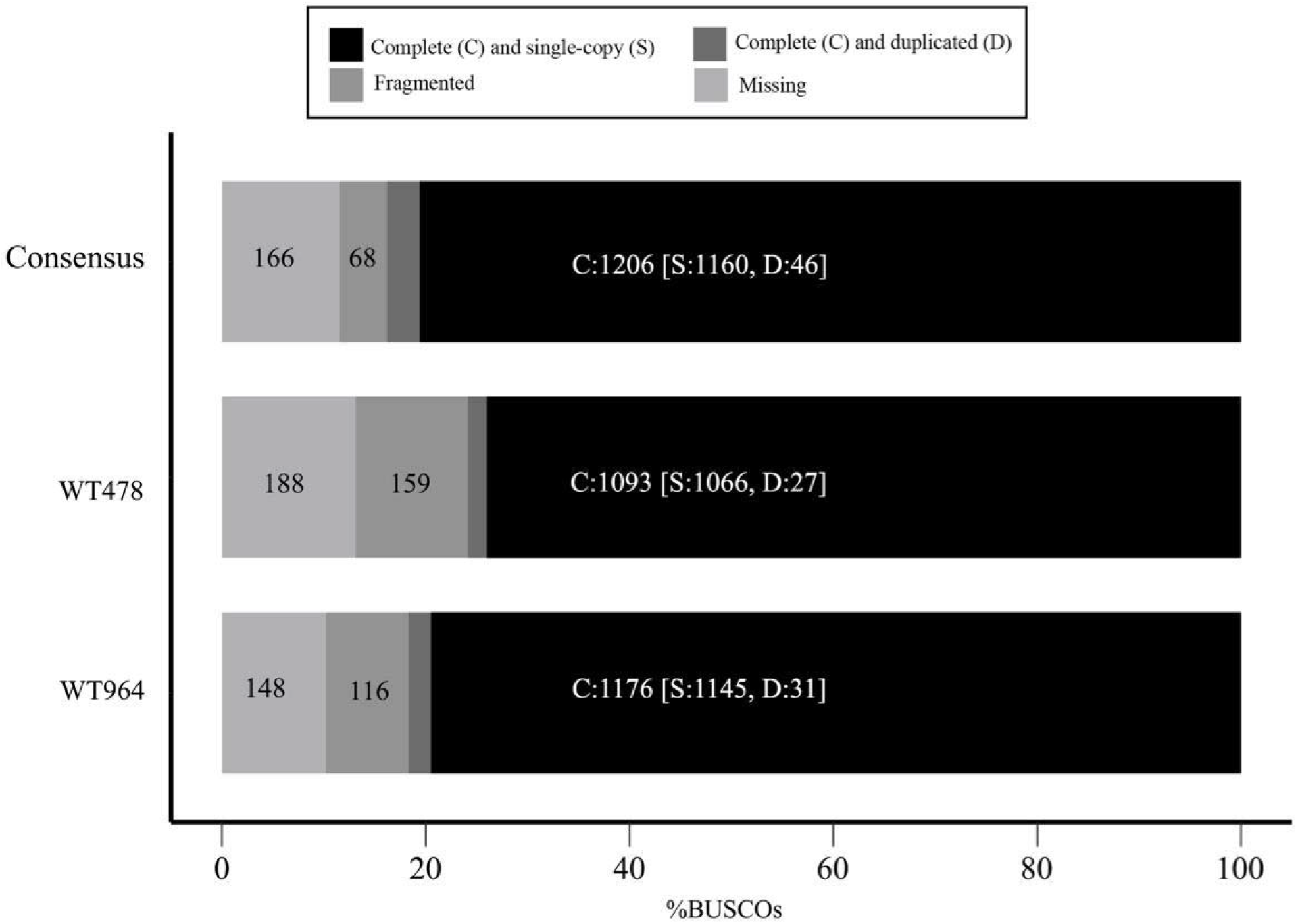
Proportion of Benchmarking Universal Single-Copy Orthologs (BUSCOs) for the *T. thalictroides* genome (total number of BUSCOs = 1,440). Data is shown for the two sequenced accessions (WT964 and WT 478) and the consensus.

Short-read data from TtWT478 had the highest level of contamination, mainly from an aphid (GCF_000142985.2) and its bacterial endosymbiont (Shigenobu and Wilson, 2011) (GCF_000009605.1) (Suppl. Fig. 1). Clean short reads were used to generate three genome assemblies (final assemblies generated with MaSuRCA v3.2.9 are reported here), one for each of the live accessions and a combined one. From here on we focus on the latter and refer to it as the “consensus assembly”, since it provides the best assembly statistics and gene coverage (Fig. 2). Mean contig coverage for the consensus assembly was 55x (median = 31.9x).

Our genome size estimate from the consensus was 243.1 Mbp, comparable to the 286.4 Mbp estimate from k-mer frequency statistics in GenomeScope (Suppl. Fig. 2) and to that of *Aquilegia coerulea*, another diploid member of the Ranunculaceae (291.7 Mb, Filiault et al., 2018). Our genome size estimate was smaller than previous values from flow cytometry (1C value=0.366 pg or 356 Mbp, Soza et al., 2013), likely due to complex or long repetitive regions (longer than the short read length) that were not recovered in our sequencing and/or assembly. Analysis of repetitive sequences with RepeatMasker and RepeatModeler, revealed that over a third of the genome (37.59%) could be assigned to different classes of repeat elements, the most abundant class being the Autonomous Long Terminal Repeat (LTR) retroelements, particularly from the Copia (7.71%) and Gypsy (6.78%) superfamilies (Suppl. Table 4). In fact, LTR transposable elements have been found to underlie homeotic flower mutants of *T. thalictroides* (Galimba et al., 2012). The GenomeScope heterozygosity estimate for the consensus assembly was 1.23% (Suppl Fig 2), which is higher than that of *A. coerulea* (0.2-0.35%, Filiault et al., 2018) but within the range of obligate outcrossers (Leffler et al., 2012).

We predicted 33,624 protein-coding genes and 1,936 non-coding RNA loci (Suppl. Table 5). Orthogroup analysis resulted in 67.4% of predicted protein-coding genes forming clusters with at least one of the four other species included in the analysis, *A. thaliana, P. somniferum, A. coerulea* and *S. lycopersicum* (Suppl. Table 6). Most of these clusters were shared by all five species, providing support for our gene predictions. TransRate analysis also showed that a large fraction of predicted proteins in *T. thalictroides* had best bi-directional BLAST hits against *A. thaliana, S. lycopersicum, A. coerulea* and *P. somniferum* (Suppl. Table 7), and 88.3% of predicted protein-coding genes had hits against InterPro member databases (https://doi.org/10.6084/m9.figshare.12302096.v1). Functional descriptions were added for 22,603 protein-coding genes using AHRD. Using the pipeline established for the development of the Plant Transcription Factor Database (PlnTFDB v3.0 (Pérez-Rodríguez et al., 2010; Riaño-Pachón et al., 2007)), we identified 1,569 protein-coding loci representing TAPs: 1,258 of these were transcription factors (TFs) that can be assigned to 68 different TF families and the remaining 311 belonged to 29 families of other types of transcriptional regulators (oTRs) (Figure 5 and Suppl. Table 8). The number of transcription factor families detected compares to that found in the genomes of *A. thaliana, S. lycopersicum, A. coerulea* and *P. somniferum* (https://doi.org/10.6084/m9.figshare.12302096.v1). The ratio of TF to oTRs was similar among the three representatives of Ranunculales (*T. thalictroides, A. coerulea* and *P. somniferum*), with approximately 4.1 TFs per oTR, and lower than estimates for *A. thaliana* and *S. lycopersicum* (5.1 TF per oTR) (Suppl. Table 8). The *T. thalictroides* genome had the lowest number of predicted TAPs per thousand loci consistent with the lower number of detected BUSCO genes (Suppl. Table 8), more likely resulting from the partial nature of the genome assembly than from an underelying biological phenomenon.

### RNA-Seq of *Thalictrum thalictroides* and *T. hernandezii* floral transcriptomes

#### Sequencing and *de novo* assembly

*De novo* transcriptome assemblies of *T. thalictroides* (Tt; GHXU00000000) consisted of 54,104 contigs (N50=1,817 bp), while *T. hernandezii* (Th; GHXT00000000) had 124,707 contigs (N50=1,703 bp), with 80.1% and 82.9% identified complete BUSCOs respectively (Table 2). Transdecoder identified 26,407 ORFs per sample for *T. thalictroides* and 52,313 for *T. hernandezii* (https://doi.org/10.6084/m9.figshare.12524036.v1). Since *T. hernandezii* is a tetraploid (Soza et al., 2013), *de novo* transcriptome assembly was expected to include up to four expressed alleles per gene, thus explaining the larger number of assembled transcripts for this species. There were approximately twice as many reciprocal best hits (CRBB) in *T. hernandezii* compared to *T. thalictroides* with either of the reference transcriptomes. Polyploidy also explains the low fraction of reads mapping back to the assembly (Table 2).

**TABLE 2.**
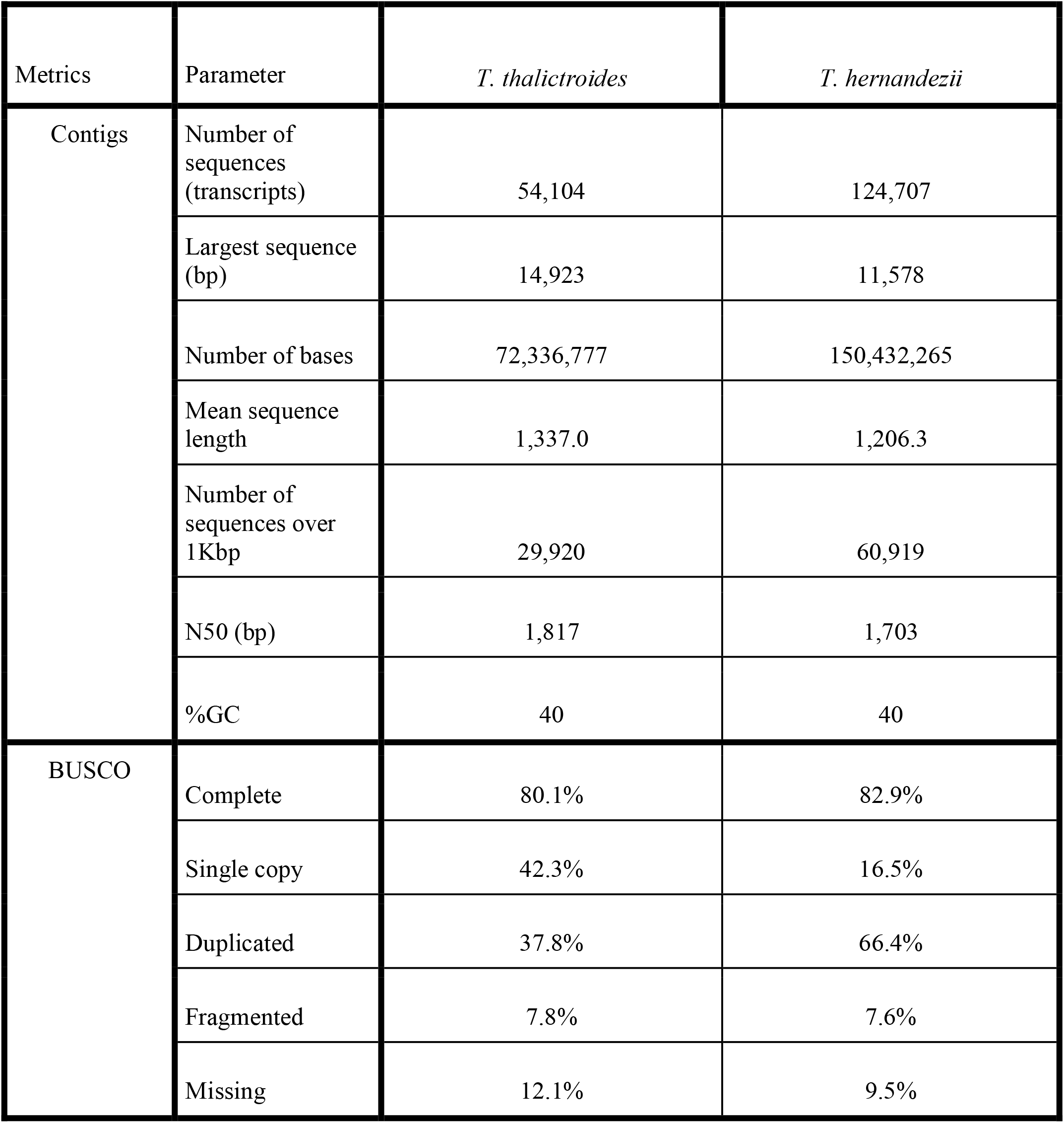
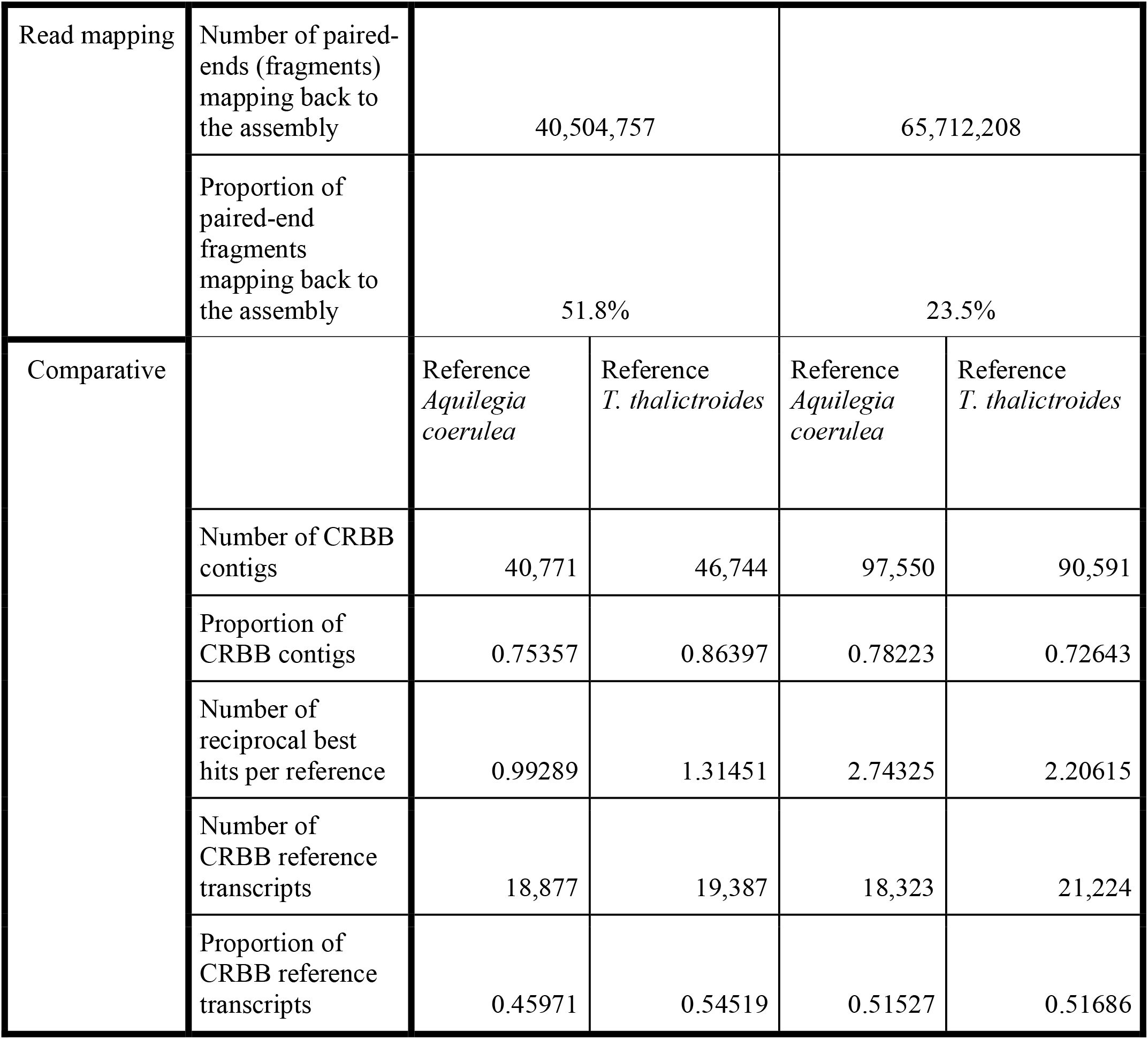
Properties of *de novo* floral transcriptome assemblies of *Thalictrum thalictroides* and *T. hernandezii*. CRBB: high-confidence predicted homologs identified by the Conditional Reciprocal Best BLAST algorithm (Aubry et al., 2014), implemented in TransRate (Smith-Unna et al., 2016).

| Metrics | Parameter | *T. thalictroides* | *T. hernandezii* |
| --- | --- | --- | --- |
| Contigs | Number of sequences (transcripts) | 54,104 | 124,707 |
|  | Largest sequence (bp) | 14,923 | 11,578 |
|  | Number of bases | 72,336,777 | 150,432,265 |
|  | Mean sequence length | 1,337.0 | 1,206.3 |
|  | Number of sequences over 1Kbp | 29,920 | 60,919 |
|  | N50 (bp) | 1,817 | 1,703 |
|  | %GC | 40 | 40 |
| BUSCO | Complete | 80.1% | 82.9% |
|  | Single copy | 42.3% | 16.5% |
|  | Duplicated | 37.8% | 66.4% |
|  | Fragmented | 7.8% | 7.6% |
|  | Missing | 12.1% | 9.5% |

|  |  |  |  |  |  |
| --- | --- | --- | --- | --- | --- |
| Read mapping | Number of paired-ends (fragments) mapping back to the assembly | 40,504,757 |  | 65,712,208 |  |
|  | Proportion of paired-end fragments mapping back to the assembly | 51.8% |  | 23.5% |  |
| Comparative |  | Reference<br>*Aquilegia coerulea* | Reference<br>*T. thalictroides* | Reference<br>*Aquilegia coerulea* | Reference<br>*T. thalictroides* |
|  | Number of CRBB contigs | 40,771 | 46,744 | 97,550 | 90,591 |
|  | Proportion of CRBB contigs | 0.75357 | 0.86397 | 0.78223 | 0.72643 |
|  | Number of reciprocal best hits per reference | 0.99289 | 1.31451 | 2.74325 | 2.20615 |
|  | Number of CRBB reference transcripts | 18,877 | 19,387 | 18,323 | 21,224 |
|  | Proportion of CRBB reference transcripts | 0.45971 | 0.54519 | 0.51527 | 0.51686 |

#### Identification of SSRs from *de novo* floral transcriptomes

SSRs are widely distributed throughout eukaryotic genomes and transcriptomes, their high variability and co-dominant inheritance makes them useful molecular markers for population-level studies (Thiel et al., 2003). A search within the *Thalictrum* floral transcriptomes detected 30,457 SSR markers, and 65,651 in the *T. thalictroides* genome, with approximately half of the analyzed contigs containing SSRs. The most common SSR in the genome of *T. thalictroides* was dinucleotide, while in the transcriptomes it was trinucleotide followed by dinucleotide (Suppl. Table 9, https://doi.org/10.6084/m9.figshare.11984370.v8).

#### High representation of transcription factors in floral transcriptomes

3541 transcripts could be assigned into 94 Transcription Associated Protein (TAP) families in the *T. thalictroides* and *T. hernandezii de novo* assembled transcriptomes. *MYB-related, AP2-EREBO* and *bZIP* were among the top families represented, with several families expressed at significantly different levels in the three transcriptomes (Fig. 5), although a more comprehensive experiment including biological replicates will be needed to quantify gene expression.

#### Functional annotation of floral transcriptomes

In order to place as many transcripts as possible within orthogroups, both *Thalictrum* transcriptomes were compared against multiple reference genomes: 1) The *Thalictrum thalictroides* draft genome (this work); 2) The model plant *Arabidopsis thaliana* (Brassicaceae), and 3) The two phylogenetically most closely-related genomes, *Aquilegia coerulea* (Ranunculaceae) and *Papaver somniferum* (Papaveraceae, Ranunculales). The use of multiple reference genomes captured most orthologs, aiding in the validation of our gene predictions and resulting in 14 orthogroups specific to *T. thalictroides* and 75 to *T. hernandezii* RNA datasets (Suppl Fig 3). Specific orthogroups could potentially result from underlying biological processes such as lineage-specific expansions, or could partly consequence of technical limitations of our datasets.

Stepwise comparisons were conducted to, a) validate transcripts against the *T. thalictroides* draft genome and b) conduct inter- and intraspecies comparisons between wind vs. insect-pollinated and male vs. hermaphrodite floral morphologies (Fig. 2). First, we performed an inter-species transcriptome comparison, using the reference genome for validation (Fig 3i). The three-way intersection in the Venn diagram (5,204 orthogroups) represents orthologs that can be mapped to the *T. thalictroides* draft reference genome and that are expressed in floral transcriptomes of both species. A “core” of 9,556 orthogroups common to both species is represented by three of the intersecting areas (5,204 + 1,298 + 3,054). The intersection area marked as “A” (3,477 orthogroups) comprises 6,451 transcripts uniquely-expressed in *T. thalictroides* that can be mapped back to the reference genome and can be therefore considered of high confidence. The intersection area marked as “B” (3,054 orthogroups), comprises 11,251 transcripts found exclusively in *T. hernandezii* and similarly validated by the reference genome. Two sets of species-specific orthogroups not found in the genome (274 and 473 orthogroups each) deserve further exploration, some of them could be due to shortcomings in our genome sequencing, yet others could represent interesting underlying processes, such as lineage-specific expansions and/or losses. As expected, a substantial number of genomic orthogroups (3,327) are not expressed, and hence not found in the floral transcriptomes. Finally, orthogroups found in both transcriptomes but not in the genome (1,298) could evidence the limitations of our draft genome, especially since most are shared with other species (Suppl. Fig. 3).

**FIGURE 3.**
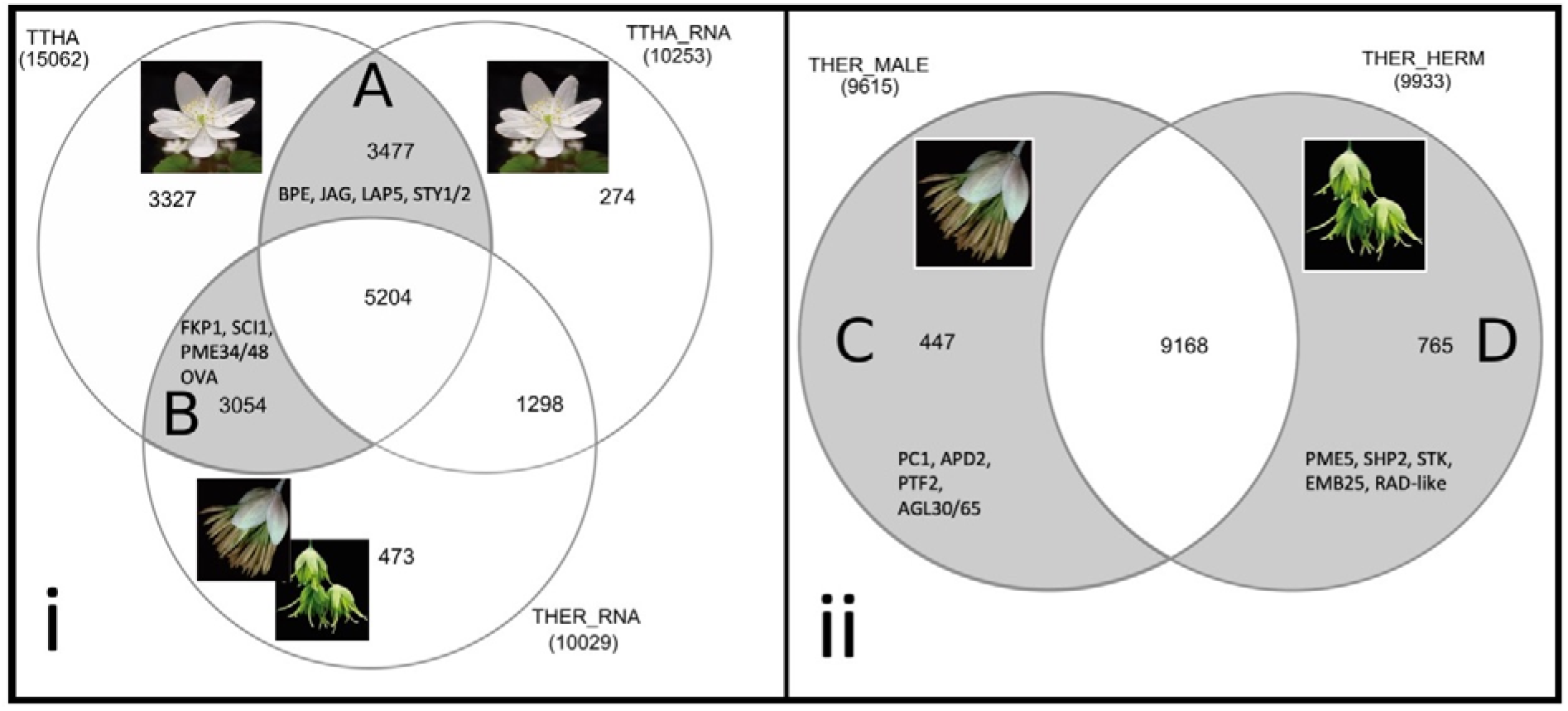
Number of shared and unique orthogroups or transcripts among the *Thalictrum* datasets in this study, with examples of candidate genes (Arabidopsis protein names used). The total number of orthogroups per dataset is indicated in parenthesis. *T. thalictroides* (Tt), insect-pollinated with hermaphrodite flowers. *T. hernandezii* (Th), wind-pollinated with hermaphrodite (Th_H) and staminate (Th_S) flowers. Representative flower pictures are shown. (i) Interspecies comparison of floral transcriptomes (Tt RNA and Th RNA) and validation against the *T. thalictroides* draft reference genome (Tt). Area “A”: *T. thalictroides-exclusive* orthogroups found in the reference genome; “B”: *T. hernandezii*-exclusive orthogroups (both floral types combined) found in the draft reference genome. (ii) Intraspecies comparison of transcripts of two floral types (male and hermaphrodite) within *T. hernandezii*. Area “C”: male flower-specific transcripts; Area “D”: hermaphrodite flower-specific transcripts.

Second, we performed an intraspecific comparison within *T. hernandezii*-specific orthogroups validated by our reference genome (Fig. 3i area B; 3,054 orthogroups). Transcripts from male (Ther_S) and hermaphrodite (Ther_H) floral transcriptomes were compared, yielding 447 male-specific and 765 hermaphrodite-specific transcripts (Fig. 3ii, areas C and D, respectively).

A GO term enrichment analysis provided a broad characterization of gene functions at the intersection between the three datasets (Fig. 4 and Suppl. Table 10), focusing on the ontology domain Biological Process. Enriched GO terms were dominated by regulatory processes (Fig. 4), and relevant categories included Gene silencing by miRNA, Maintenance of floral organ identity, Meristem initiation, Adaxial/abaxial pattern specification and Negative regulation of growth, amongst others (Suppl. Table 10, https://doi.org/10.6084/m9.figshare.12465254.v1).

**FIGURE 4.**
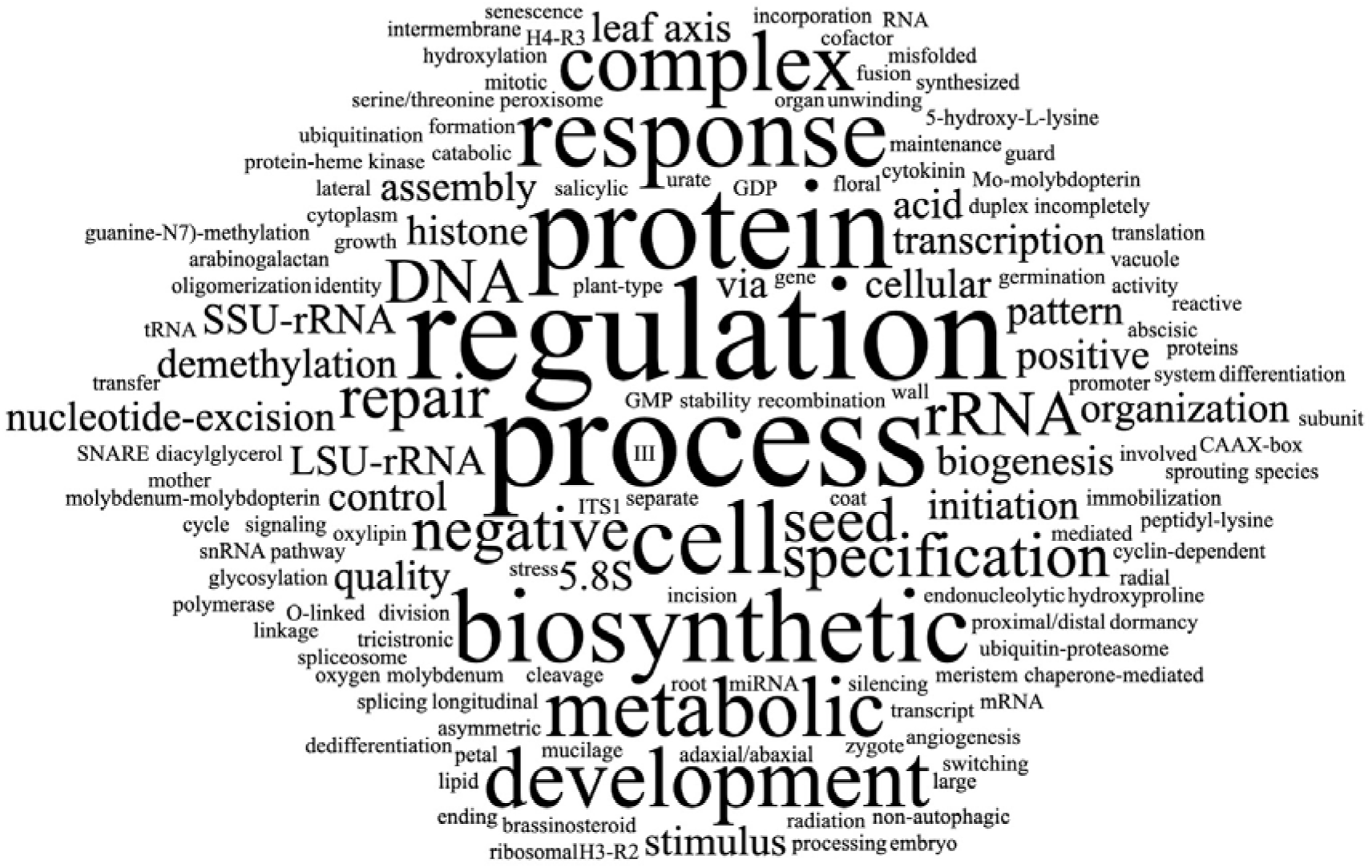
Word cloud for Enriched Gene Ontology (GO) categories among floral transcriptomes of two distinct *Thalictrum* species, emphasizing the predominant role of regulatory processes. GO terms for Biological Process (P-adj < 0.05) correspond to *T. hernandezii*-specific orthogroups not expressed in *T. thalictroides*, and validated against the reference genome (see Fig. 2i, area B). A complete list of enriched terms can be found in Suppl. Table 8.

**FIGURE 5.**
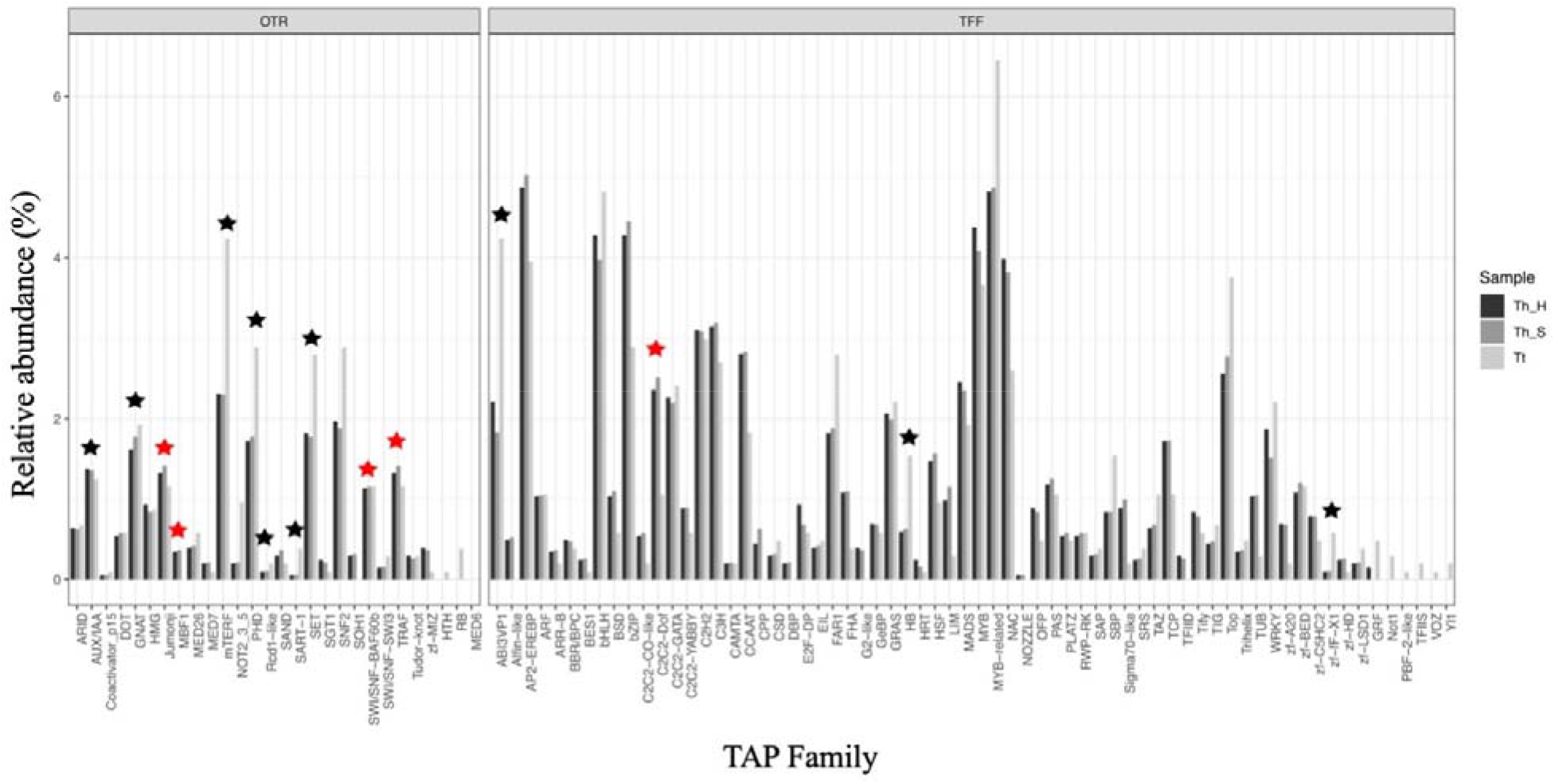
The relative abundance of different families of Transcription Associated Proteins (TAP) in the floral transcriptomes of *Thalictrum sp. T. thalictroides* (Tt); *T. hernandezii* (Th_H, hermaphrodite and Th_S, staminate) floral transcriptomes. Transcripts above 0.1 TPM (transcripts per million) included. Statistically significant differences between samples (Chi-Square test) are marked with black (P < 0.01) or red (P < 0.05) asterisks. TFF: Transcription Factor Family, OTR: Other Transcriptional Regulator Family (after Riaño-Pachón et al., 2007).

#### Data-mining examples: Potential candidate genes

To test applications of our results, we identified potential candidates for the morphological differences between flower types in the two species (Fig. 1). Our goal was to provide a preliminary, qualitative working list of candidate genes for future investigations of the genetic basis of distinct sexual systems (hermaphroditic or unisexual) and pollination modes (insect or wind). To that end, we first searched for known candidate genes within our comparisons (Fig. 3 and section above).

Venn diagram areas A and B (Fig 3i) represent functional orthogroups for flowers with distinct morphologies due to differing pollination modes (insect-pollinated *T. thalictroides* vs. wind-pollinated *T. hernandezii*). Venn diagram areas C and D (Fig. 3ii) represent examples of transcripts expressed in flowers with distinct sexual systems: staminate flowers, with sepals and stamens; hermaphrodite flowers, with added carpels. *T. thalictroides* has petaloid sepals that are comparatively bigger and white, upright stamens with smaller anthers and carpels with short styles and stigmas. *T. hernandezii* has smaller flowers with smaller, green sepals, pendant stamens with larger anthers on longer filaments, and carpels with longer styles and stigmas (Fig. 1 and Di Stilio et al., 2009). Based on these phenotypic differences, we predicted that the transcriptome comparisons could yield genes involved in processes such as cell elongation or cell division (longer stamen filaments and styles), pectins (increased flexibility in pendant filaments), epidermal cell elongation (extended stigmatic papillae in wind-pollinated flowers), or increased pollinator grip in petaloid organs (Di Stilio et al., 2009), amongst others. A subset of the genes emerging from our comparisons fit these criteria, thus serving as validation for the usefulness of our datasets as a resource (see below).

First, we searched for previously-characterized candidate genes in the *T. thalictroides* reference genome and the three transcriptomes. A homolog of *MIXTA-like2* (*ThtMYBML2*, FJ487606.1) with a role in papillate cells and stigmatic papillae (Di Stilio et al., 2009) was identified in both species, as a full transcript in the genome assembly (GenBank: KAF5204412.1) and the Th transcriptome (90% protein identity, TSA:GHXT01017115) and in two fragments in the Tt transcriptome assembly (TSA:GHXU01051721 and GHXU01036124). A second candidate for differences in morphology between species is the *Thalictrum STYLE2.1* ortholog, involved in style length in tomato (Chen et al., 2007). A *Solanum* query (Uniprot B6CG44) was used to retrieve sequences from the *T. thalictroides* genome (65% protein identity, GenBank: KAF5189420.1), and the transcriptomes (65% identity, TSA:GHXU01012872 and 67% identity, TSA:GHXT01076288). The presence of these two previously-characterized candidate genes validated the usefulness of our genome and transcriptome assemblies for obtaining full-length sequences of genes of interest. Their presence at the three way intersection of all datasets (Fig. 3i) suggests that regulatory changes in expression levels, rather than on/off switches, likely underlie the phenotype differences.

We then proceeded to perform a transcription factor (TF) family enrichment analysis for the areas of interest (areas A-D, Fig. 3 and Suppl. Table 11). An example of a relevant enriched TF families for *T. thalictroides* (area A, Fig. 3i) included bHLH, involved in jasmonic acid signaling that affects stamen development (filament elongation), anthocyanin production (present in pink sepals) and trichome development (found on sepals) (Chen et al., 2016; Goossens et al., 2017). For *T. hernandezii* (area B, Fig. 3i) enriched TF families included: 1) C2C2 GATA that are regulated by the B class flower organ identity genes (Zik and Irish, 2003) and affect chlorophyll synthesis, relevant to green flowers (Bi et al., 2005); 2) SAP that regulate organ size in these small flowers (Wang et al., 2016) and affect ovule development (Byzova et al., 1999) and 3) C2C2 CO-like that control flowering (Steinbach, 2019). GRF family was a promising candidate for hermaphrodite flowers of *T. hernandezii* (Fig. 3ii, area D), given their role in cell proliferation leading in organ size and carpel development (Lee et al., 2018), while members of the cell proliferation pathway CPP (Yang et al., 2008) were enriched in the male flowers (Fig. 3ii area C), with their dangling stamens on long filaments. MADS-box family genes were enriched throughout, as expected given their prominent role in all aspects of flower development.

Finally, we identified candidate orthogroups or transcripts of interest in our dataset via stepwise comparisons (https://doi.org/10.6084/m9.figshare.11984358.v3). Orthogroups exclusive to *T. thalictroides* (Fig. 3i, area A) and relevant to insect pollination syndrome included orthologs of the Arabidopsis proteins BIG PETAL (BPE, Szécsi et al., 2006) and JAGGED (JAG, Sauret-Güeto et al., 2013) with potential role in the large, petaloid sepals (Fig. 1A), LESS ADHESIVE POLLEN 5 (LAP5, Dobritsa et al., 2010) in regulating the degree of pollen stickiness (higher in insect-pollinated species), STYLISH 1/2 (STY1/2, Sohlberg et al., 2006) in style length (short) and stigma size (compact or “capitate”), and jasmonate (JA) hormone pathway JAZ 1-3 (Browse and Wallis, 2019; Figueroa and Browse, 2015) in stamen filament elongation (shorter and flattened). Other usual suspects in flower development enriched in this area included WUSCHEL-related homeoBOX (WOX3-4) that contribute to general aspects of floral architecture and morphology (Costanzo et al., 2014), the YABBY protein INNER-NO-OUTER (INO) in ovule development (Simon et al., 2017) and the TCP protein CYCLOIDEA (CYC) in flower symmetry and general flower form (Hileman and Cubas, 2009).

*T. hernandezii*-specific genes relevant to wind pollination syndrome (Fig. 3i, area B) included orhtologs of Arabidopsis FLAKY POLLEN 1 (FKP1) affecting pollen coat qualities (Ishiguro et al., 2010) and thus relevant to pollen adaptations to wind pollination; the PECTIN METHYLESTERASES 34 (PM34) highly expressed in stamen filaments and relevant to the overly elongated linear filaments and styles (Gou et al., 2012; Irshad et al., 2008) and PME48, specific to pollen tube elongation and thus relevant for successful fertilization through the long styles (Leroux et al., 2015); the stigma and style cell-cycle inhibitor 1 (SCI1, DePaoli et al., 2014), relevant to the extended styles and stigmas found in this species; and most members of the OVULE ABORTION (OVA) family (Berg et al., 2005), which could play a role in the differential floral sexes.

Comparisons between *T. hernandezii* hermaphrodite and staminate flower transcriptomes (intraindividual, Fig. 3 ii) were best suited to identify carpel-specific candidate genes (only present in hermaphrodite flowers) and, to a lesser extent, genes related to sexually dimorphic features of stamens and sepals (present in both flower types). Out of 765 transcripts uniquely expressed in *T. hernandezii* hermaphrodite flowers (Fig. 3ii, area D); relevant candidates included transcripts orthologous to PECTIN METHYL ESTERASE 5 (PME5) expressed in carpels and during fruit development (Louvet et al., 2006), SHATERPROOF 2 (SHP2) and SEEDSTICK (STK) in ovule development (Favaro et al., 2003), EMBRYO DEFECTIVE 25 (EMB25) in embryo development (Meinke, 2020) and RADIALIS-like (RAD-like) proteins with roles in ovule and embryo development and potentially in flower symmetry (Baxter et al., 2007). Among the 417 male-specific transcripts, it is worth noting differential transcripts for the main contributors to pollen and pollen-tube function MIKC* MADS box genes AG-like30/65 (Adamczyk and Fernandez, 2009), and several coding for orthologs to other Arabidopsis proteins related to pollen development such as POLLEN CALCIUM BINDING PROTEIN 1 (PC1, Wang et al., 2008), ABERRANT POLLEN DEVELOPMENT 2 (APD2, Luo et al., 2012a) and POLLEN EXPRESSED TRANSCRIPTION FACTOR 2 affecting pollen germination (PTF2, Niu et al., 2013).

### Conclusions

This study provides the first *Thalictrum* genome, and transcriptomic resources, for two species that represent an early-diverging lineage of dicotyledonous plants with distinct floral morphologies related to different sexual and pollination systems. The application of these resources is exemplified in our results, which include the identification of transposable elements, molecular markers and putative candidate genes. Future uses of these resources include the identification of other genes of interest and their regulatory regions (draft genome) as well as primer design for phylogenetic and population-level studies.

## ACKNOWLEDGMENTS

This work was funded through the Research and Conference Grants Administration System (RCGAS) of The University of Hong Kong’s Small Project Funding (to TA); and grants from NSF DEB 1911539 (Opus-MCS) and The Fred Gloeckner Foundation, Inc. (to VSD). Access to computational resources was granted through the University of Washington, USA, the Earlham Institute, UK, Institut National de la Recherche Agronomique (INRA) and the Center for Nuclear Energy in Agriculture, University of São Paulo, Brazil. DMRP is a level 2 CNPq research fellow – Brazil (310080/2018-5)

We thank Gemy George Kaithakottil from the Earlham Institute UK, Eric Montaudon from the Institut National de la Recherche Agronomique (INRA), Jorge Mario Muñoz and Juliana Arcila from the Comparative Biology Laboratory at La Corporación para Investigaciones Biológicas (CIB) for help with scripts for this work. Luis Eduardo Mejia for assistance with figures, Diego Morales-Briones for assistance with SRA submission and Valerie Soza for technical assistance in the laboratory.

RepeatModeler Download Page.

