## Supplementary tables for "Facilitating candidate gene discovery in an emerging model plant lineage: Transcriptomic and genomic resources for *Thalictrum* (Ranunculaceae)"

**SUPPLEMENTARY FIGURES AND TABLES**

Suppl. figure 1. Phylum-level taxonomic assignment of contaminants in the genome of *Thalictrum thalictroides* (obtained with BlobPlots). The top plot summarizes sequence coverage and GC content for each contig in the assembly (circles), with *T. thalictroides* contigs shown in yellow. The size of the circles is proportional to contig size; colors indicate taxonomic assignment, as shown in the inset. The bottom plot shows the frequency distribution of the contaminating taxa for each of the two accessions used.

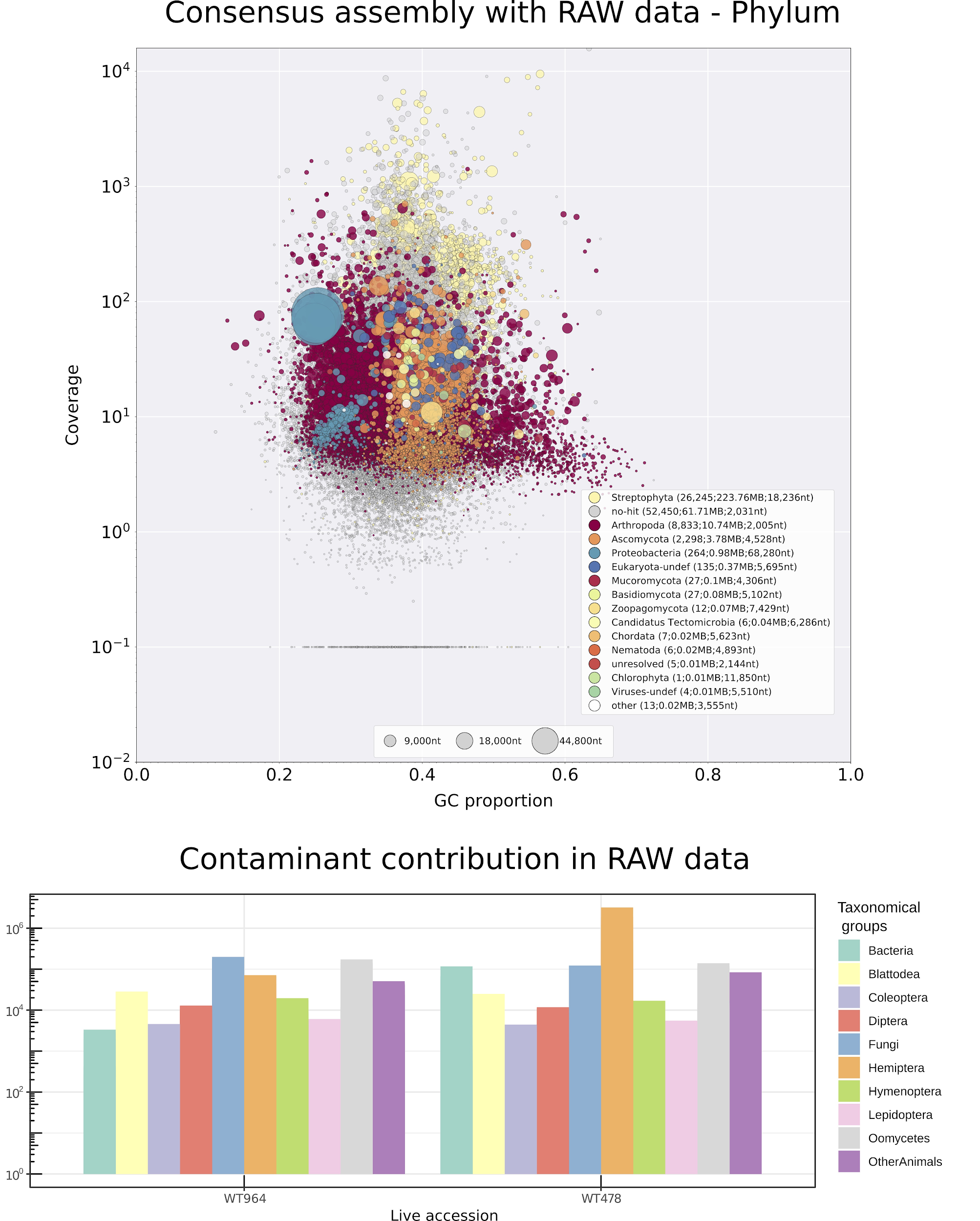

Suppl. Figure 2. Genome size estimation with GenomeScope. len: inferred total genome length, uniq: percent of the genome that is unique (not repetitive); het: overall rate of heterozygosity; kcov: mean kmer coverage for heterozygous bases; err: error rate of the reads; dup: average rate of read duplications; k:17: size of k-mer to perform the analysis; cov-threshold: maximum number of k-mer coverage taken into account for analysis, kmer with higher coverage could be organelles or contaminants.

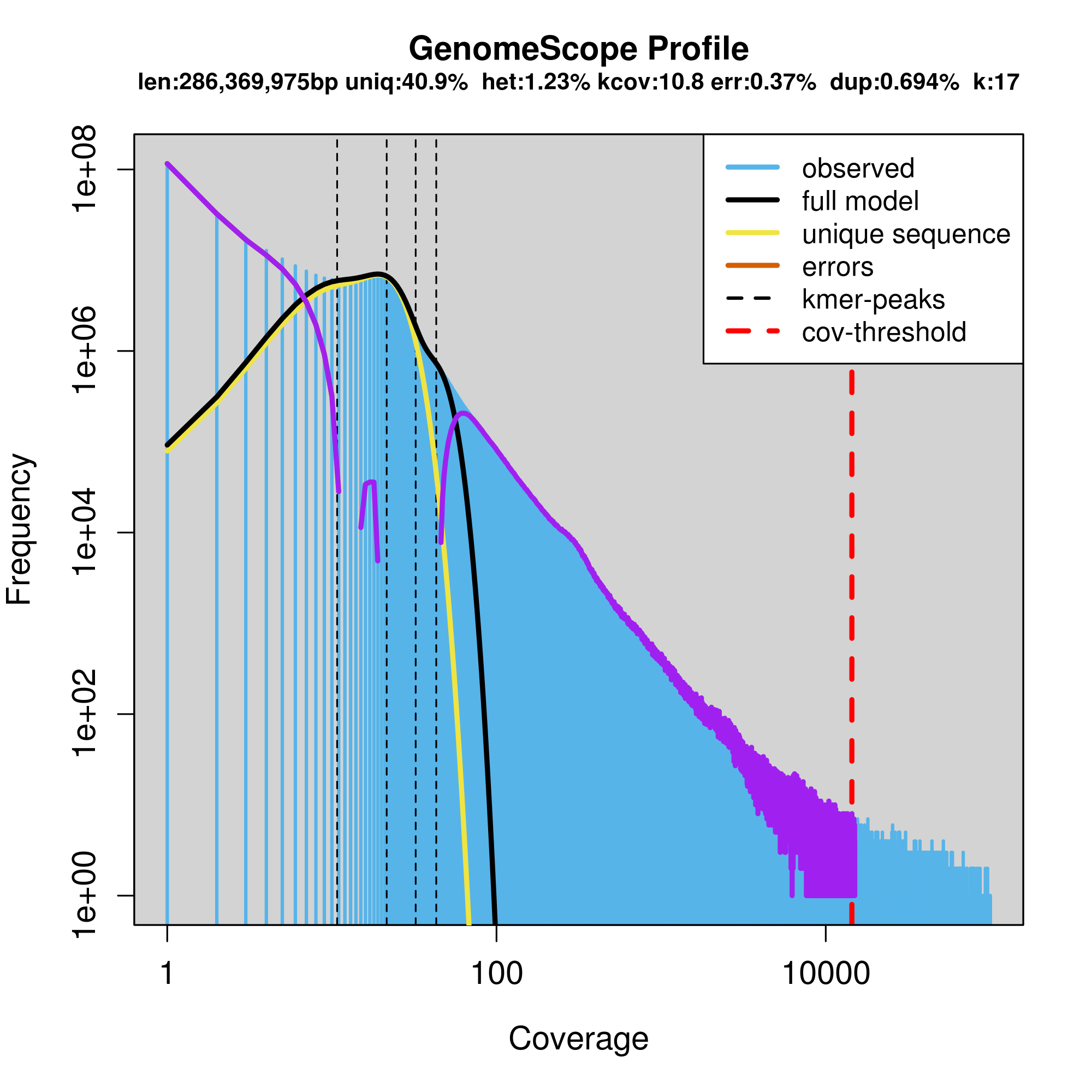

Suppl. figure 3. The number of shared and unique orthologous genes (orthogroups) at the intersection between five species and six datasets. Genome comparisons between *Thalictrum thalictroides* (TTHA, this study) and the reference genomes of *Arabidopsis thaliana* (ATHA), *Aquilegia coerulea* (ACOE) and *Papaver somniferum* (PSOM); and two *Thalictrum* floral transcriptomes, *T. thalictroides* (TTHA_RNA) and *T. hernandezii* (THER_RNA; this study).

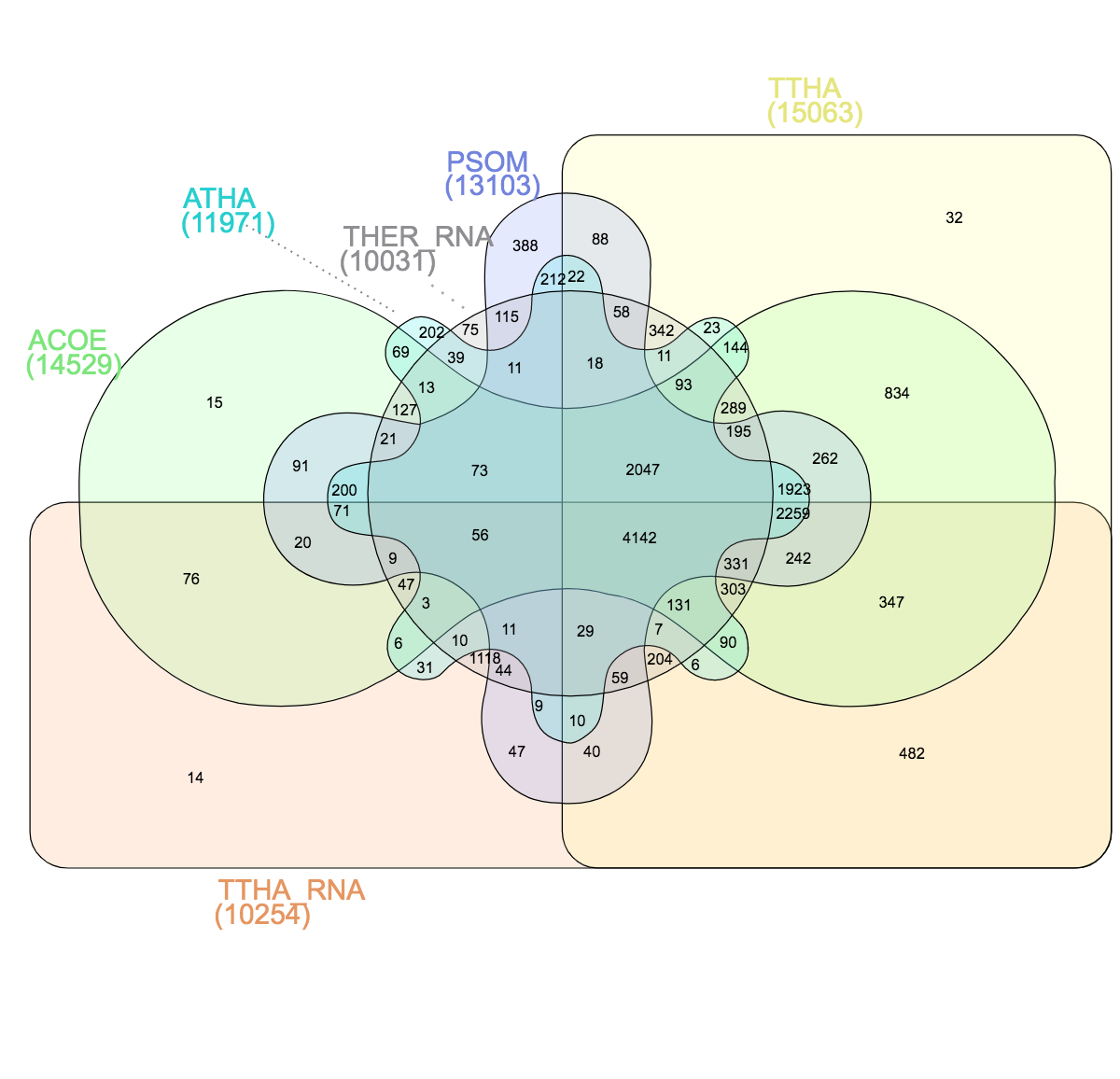

Suppl. Table 1. Voucher and source information for species of *Thalictrum* in this study.

| **Species** | | **DNA code** | **Usage** | **Voucher accession no. (Herbarium)** | **Locality** | **SRA Accession Number** |
| --- | --- | --- | --- | --- | --- | --- |
| *T. hernandezii* | ThWT441 | | RNA-seq analysis | V. Di Stilio 119 (WTU) | Cultivated from wild seed, Huitzila, Mexico Liston1215 | SRR6869420, SRR6869419 |
| *T .thalictroides* | TtWT478 | | Whole genome assembly and RNA-seq analysis | V. Di Stilio 124 (WTU) | Cultivated from nursery material (MI, USA) | Genome: SRR6869426, SRR6869425, SRR6869418  Transcriptome:  [SRR6869422](https://trace.ncbi.nlm.nih.gov/Traces/sra/?run=SRR6869422)  GBVZ |
| *T. thalictroides* | TtWT964 | | Whole genome assembly | Unvouchered | Cultivated from nursery material (NC, USA) | Genome: SRR6869424, SRR6869423, SRR6869421 |

Suppl. Table 2. Summary statistics and accession numbers for *Thalictrum thalictroides* genomic sequencing. Data for two genetically distinct individuals shown (WT964 and WT478) and three NGS libraries of different insert sizes. Trimming involved contaminant removal for improved sequence quality (see Suppl. Fig. 1).

| Sample | Library insert size | Number of sequenced fragments | | | | | SRA Accession number |
| --- | --- | --- | --- | --- | --- | --- | --- |
|  |  | Raw Reads | Trimmed reads | Percent (from Raw) after Trimming | Clean reads | Percent (from Raw) after Cleaning |  |
| WT964 | 170 | 11,429,928 | 10,146,420 | 88.8 | 9,758,045 | 85.4 | SRX3823744 |
| WT964 | 500 | 11,887,778 | 9,155,669 | 77.0 | 8,910,174 | 75.0 | SRX3823741 |
| WT964 | 800 | 11,826,149 | 9,253,079 | 78.2 | 9,130,960 | 77.2 | SRX3823742 |
| WT478 | 170 | 11,484,802 | 10,191,821 | 88.7 | 6,601,641 | 57.5 | SRX3823739 |
| WT478 | 500 | 11,477,882 | 10,414,934 | 90.7 | 7,727,941 | 67.3 | SRX3823740 |
| WT478 | 800 | 11,721,131 | 9,012,622 | 76.9 | 6,977,136 | 59.5 | SRX3823747 |

Suppl. Table 3. Frequency of contig length in *de novo* genome assemblies of *Thalictrum thalictroides*. Data shown is for two genetically distinct individuals (WT964 and WT478) and the consensus.

|  | Contig Size | Consensus | WT964 | WT478 |
| --- | --- | --- | --- | --- |
| Number of contigs | >= 5000 bp | 13,086 | 12,064 | 10,069 |
|  | >= 10000 bp | 6,055 | 4,878 | 3,157 |
|  | >= 25000 bp | 1,312 | 644 | 171 |
|  | >= 50000 bp | 154 | 35 | 5 |

Suppl. Table 4. Transposable and other repeat elements in the *Thalictrum thalictroides* genome.

| **Class** | **Count** | **Number of bp** | **% of genome** |
| --- | --- | --- | --- |
| DNA | 924 | 180559 | 0.07% |
| CMC | 4 | 427 | 0.00% |
| CMC-EnSpm | 2122 | 772578 | 0.32% |
| MULE-MuDR | 1405 | 264413 | 0.11% |
| MULE-NOF | 463 | 102821 | 0.04% |
| Maverick | 161 | 131215 | 0.05% |
| Merlin | 1 | 71 | 0.00% |
| MuLE-MuDR | 1649 | 943946 | 0.39% |
| Novosib | 4 | 146 | 0.00% |
| P | 78 | 26495 | 0.01% |
| PIF-Harbinger | 2067 | 782953 | 0.32% |
| Sola-1 | 7 | 704 | 0.00% |
| TcMar | 1 | 60 | 0.00% |
| TcMar-Pogo | 377 | 55449 | 0.02% |
| TcMar-Stowaway | 433 | 61511 | 0.03% |
| hAT | 1199 | 224720 | 0.09% |
| hAT-Ac | 6579 | 1106052 | 0.45% |
| hAT-Charlie | 240 | 70775 | 0.03% |
| hAT-Tag1 | 1992 | 455673 | 0.19% |
| hAT-Tip100 | 429 | 191141 | 0.08% |
| LINE | 1 | 65 | 0.00% |
| CRE | 151 | 67620 | 0.03% |
| CRE-II | 1126 | 297400 | 0.12% |
| I | 51 | 41760 | 0.02% |
| L1 | 10444 | 5198061 | 2.14% |
| L1-Tx1 | 1 | 94 | 0.00% |
| L2 | 165 | 115048 | 0.05% |
| Penelope | 86 | 32929 | 0.01% |
| RTE-BovB | 171 | 38016 | 0.02% |
| RTE-X | 1 | 64 | 0.00% |
| LTR | 1188 | 125379 | 0.05% |
| Cassandra | 1130 | 252592 | 0.10% |
| Caulimovirus | 209 | 80928 | 0.03% |
| Copia | 23671 | 18742363 | 7.71% |
| DIRS | 1 | 59 | 0.00% |
| ERV1 | 51 | 18648 | 0.01% |
| Gypsy | 19286 | 16470704 | 6.78% |
| RC | -- | -- | -- |
| Helitron | 1541 | 920228 | 0.38% |
| Helitron-2 | 2 | 143 | 0.00% |
| Retroposon | -- | -- | -- |
| L1-derived | 1 | 130 | 0.00% |
| SINE | 36 | 3579 | 0.00% |
| ID | 31 | 2576 | 0.00% |
| tRNA-RTE | 3 | 274 | 0.00% |
| Unknown | 147569 | 41841755 | 17.21% |
| **total interspersed** | **227051** | **89622124** | **36.87%** |
| Low_complexity | 7710 | 368028 | 0.15% |
| Satellite | 101 | 18244 | 0.01% |
| Simple_repeat | 31601 | 1369568 | 0.56% |
| **Total** | **266463** | **91377964** | **37.59%** |

Suppl. Table 5 Number of predicted RNA gene types in the *T. thalictroides* genome.

| **Gene Type** | **Number of Genomic Loci** |
| --- | --- |
| mRNA | 33,624 |
| snoRNA | 1,125 |
| tRNA | 557 |
| snRNA | 153 |
| miRNA | 74 |
| SRP RNA | 13 |
| Autocatalytically-spliced intron | 8 |
| RNase MRP RNA | 2 |
| rRNA | 2 |
| ribozyme | 1 |
| scRNA | 1 |
| TOTAL | 35,560 |

Suppl. Table 6. Identification of orthogroups in the *T. thalictroides* genome, using columbine, tomato, opium poppy and Arabidopsis. Protein identifiers for each orthogroup and species are available at https://doi.org/10.6084/m9.figshare.11984358.v1

| **Species** | **Species abbreviation** | **Number of Genes in Orthogroups** | **Total Number of Genes** | **Percent of genes assigned to orthogroups** | **Number of Orthogroups** |
| --- | --- | --- | --- | --- | --- |
| *Thalictrum thalictroides* | TTHA | 22671 | 33626 | 67.4 | 14385 |
| *Aquilegia coerulea* | ACOE | 33311 | 41063 | 81.1 | 14384 |
| *Solanum lycopersicum* | SLYC | 22870 | 35768 | 63.9 | 13113 |
| *Papaver somniferum* | PSOM | 51655 | 84179 | 61.4 | 13432 |
| *Arabidopsis thaliana* | ATHA | 34190 | 48359 | 70.7 | 12593 |

Suppl. table 7. Evaluation of the predicted transcriptome from the *T. thalictroides* genome assembly (from TransRate) against four reference genomes (from Suppl. Table 6). Conditional Reciprocal Best BLAST (CRBBs) are high-confidence predicted homologs (Aubry et al., 2014; Smith-Unna et al., 2016).

| **Metric (with CRBB)** | **Reference genomes** | | | |
| --- | --- | --- | --- | --- |
|  | **ACOE** | **PSOM** | **ATHA** | **SLYC** |
| Number of Tt transcripts | 20,536 | 17,134 | 16,212 | 16,695 |
| Fraction of Tt transcripts | 0.61 | 0.51 | 0.48 | 0.5 |
| Number of reference transcripts | 17,474 | 14,075 | 12,688 | 12,674 |
| Fraction of reference transcripts | 0.43 | 0.17 | 0.26 | 0.35 |

Suppl. table 8. The number and type of Transcription Associated Proteins (TAPs) and proportion of complete BUSCO in the genomes of *T. thalictroides* (TTHA), *A. coerulea* (ACOE), *P. somniferum* (PSOM), *Solanum lycopersicum* (SLYC) and *A. thaliana* (ATHA) . TF: transcription factors; oTR: other types of transcriptional regulators.

| **Species** | **Complete BUSCOs** | **Type of TAP** | **Number of TAP loci** | **Per thousand TAP loci** | **Total number of loci** | **Ratio TF/oTR** |
| --- | --- | --- | --- | --- | --- | --- |
| ACOE | 95.8% | OTR | 296 | 11.9 | 24,823 | 4.23 |
| ACOE |  | TFF | 1,252 | 50.4 | 24,823 |  |
| ATHA | 99.6% | OTR | 342 | 12.4 | 27,655 | 5.13 |
| ATHA |  | TFF | 1,753 | 63.4 | 27,655 |  |
| PSOM | 95.5% | OTR | 743 | 11.8 | 63,012 | 4.13 |
| PSOM |  | TFF | 3,066 | 48.7 | 63,012 |  |
| SLYC | 94.5% | OTR | 388 | 10.8 | 35,768 | 5.18 |
| SLYC |  | TFF | 2,011 | 56.2 | 35,768 |  |
| TTHA | 84.5% | OTR | 311 | 9.2 | 33,626 | 4.05 |
| TTHA |  | TFF | 1,258 | 37.4 | 33,626 |  |

Suppl, table 9. Simple Sequence Repeat (SSR) motif distribution in floral transcriptomes of *T. thalictroides* and *T. hernandezii* (hermaphrodite and staminate combined) and genome of *T. thalictroides.* Abundance at or above 5% of total SSRs in the genome.

SSRs available at https://doi.org/10.6084/m9.figshare.11984370.v8

|  | *T. thalictroides* genome | *T. thalictroides*  Transcriptome | *T. hernandezii*  Transcriptome |
| --- | --- | --- | --- |
| Total Number of identified SSRs | 65,651 | 11,826 | 18,631 |
| No of SSRs present in tandem | 3,582 | 381 | 538 |
| No of contigs with SSR | 22,867 | 9,844 | 16,512 |
| No of contigs with more than 1 SSRs | 12,596 | 1,634 | 1,898 |

Suppl. table 10 . Enriched GO categories in comparisons among floral transcriptomes of *Thalictrum thalictroides* (Tt) and *T. hernandezii* (Th) at FDR< 0.05, for Biological Process (refer to Fig. 3 for areas).

| **Venn Area**  **(Fig. 2)** | **GO term** | **Annotation** |
| --- | --- | --- |
| A  (Tt specific) | GO:0032259 | methylation |
|  | GO:0001522 | pseudouridine synthesis |
|  | GO:0046940 | nucleoside monophosphate phosphorylation |
|  | GO:0000244 | spliceosomal tri-snRNP complex assembly |
| B  (Th specific) | GO:1900056 | negative regulation of leaf senescence |
|  | GO:0010405 | arabinogalactan protein metabolic process |
|  | GO:0018258 | protein O-linked glycosylation via hydroxyproline |
|  | GO:0000079 | regulation of cyclin-dependent protein serine/threonine kinase activity |
|  | GO:0048024 | regulation of mRNA splicing, via spliceosome |
|  | GO:0006515 | protein quality control for misfolded or incompletely synthesized proteins |
|  | GO:0051259 | protein complex oligomerization |
|  | GO:0006777 | Mo-molybdopterin cofactor biosynthetic process |
|  | GO:0071629 | cytoplasm protein quality control by the ubiquitin-proteasome system |
|  | GO:0070078 | histone H3-R2 demethylation |
|  | GO:0070079 | histone H4-R3 demethylation |
|  | GO:0071586 | CAAX-box protein processing |
|  | GO:0018395 | peptidyl-lysine hydroxylation to 5-hydroxy-L-lysine |
|  | GO:0002040 | sprouting angiogenesis |
|  | GO:0009688 | abscisic acid biosynthetic process |
|  | GO:0048441 | petal development |
|  | GO:0042273 | ribosomal large subunit biogenesis |
|  | GO:0006412 | translation |
|  | GO:0120009 | intermembrane lipid transfer |
|  | GO:2000377 | regulation of reactive oxygen species metabolic process |
|  | GO:0009793 | embryo development ending in seed dormancy |
|  | GO:0017003 | protein-heme linkage |
|  | GO:0106004 | tRNA (guanine-N7)-methylation |
|  | GO:0035542 | regulation of SNARE complex assembly |
|  | GO:0016567 | protein ubiquitination |
|  | GO:0051131 | chaperone-mediated protein complex assembly |
|  | GO:0010444 | guard mother cell differentiation |
|  | GO:1901333 | positive regulation of lateral root development |
|  | GO:0032889 | regulation of vacuole fusion, non-autophagic |
|  | GO:0043697 | cell dedifferentiation |
|  | GO:0009945 | radial axis specification |
|  | GO:0060184 | cell cycle switching |
|  | GO:0009301 | snRNA transcription |
|  | GO:0006384 | transcription initiation from RNA polymerase III promoter |
|  | GO:0071368 | cellular response to cytokinin stimulus |
|  | GO:0009942 | longitudinal axis specification |
|  | GO:0006281 | DNA repair |
|  | GO:0031408 | oxylipin biosynthetic process |
|  | GO:0007031 | peroxisome organization |
|  | GO:0046037 | GMP metabolic process |
|  | GO:0033683 | nucleotide-excision repair, DNA incision |
|  | GO:0046710 | GDP metabolic process |
|  | GO:0071446 | cellular response to salicylic acid stimulus |
|  | GO:0000447 | endonucleolytic cleavage in ITS1 to separate SSU-rRNA from 5.8S rRNA and LSU-rRNA from tricistronic rRNA transcript (SSU-rRNA, 5.8S rRNA, LSU-rRNA) |
|  | GO:1900458 | negative regulation of brassinosteroid mediated signaling pathway |
|  | GO:0045926 | negative regulation of growth |
|  | GO:0010014 | meristem initiation |
|  | GO:0009314 | response to radiation |
|  | GO:0009411 | response to UV |
|  | GO:0048497 | maintenance of floral organ identity |
|  | GO:0006651 | diacylglycerol biosynthetic process |
|  | GO:0035902 | response to immobilization stress |
|  | GO:0018315 | molybdenum incorporation into molybdenum-molybdopterin complex |
|  | GO:0019628 | urate catabolic process |
|  | GO:0000717 | nucleotide-excision repair, DNA duplex unwinding |
|  | GO:0010589 | leaf proximal/distal pattern formation |
|  | GO:0010070 | zygote asymmetric cell division |
|  | GO:0045951 | positive regulation of mitotic recombination |
|  | GO:0061635 | regulation of protein complex stability |
|  | GO:0071669 | plant-type cell wall organization or biogenesis |
|  | GO:0009955 | adaxial/abaxial pattern specification |
|  | GO:0048354 | mucilage biosynthetic process involved in seed coat development |
|  | GO:0009845 | seed germination |
|  | GO:0035195 | gene silencing by miRNA |

Suppl. table 11. Enriched Transcription Factor families in *Thalictrum* floral transcriptomes. Comparisons between *T. thalictroides* (Tt) and *T. hernandezii* (Th) floral transcriptomes (male flowers, Th-S and hermaphrodite flowers, Th_H). Refer to Fig. 3 for “Area”.

FDR=False Discovery Rate.

| **Area** | **Transcription Factor Families** | **FDR** |
| --- | --- | --- |
| A  (Tt specific) | mTERF | <0.05 |
|  | bHLH | <0.1 |
|  | FAR1 |  |
|  | MADS-box |  |
| B  (Th specific) | C2C2-CO-like | <0.05 |
|  | MYB-related | <0.1 |
|  | SNF2 |  |
|  | DDT |  |
|  | C2C2-GATA |  |
|  | SWI/SNF-BAF60b |  |
|  | CCAAT |  |
|  | mTERF |  |
|  | CPP |  |
|  | Alfin-like |  |
|  | MADS-box |  |
|  | PAS |  |
|  | SAP |  |
|  | DBP |  |
|  | FHA |  |
|  | zf-MIZ |  |
|  | C3H |  |
|  | C2C2-Dof |  |
|  | TRAF |  |
| C  (Th_S) | CCP | <0.05 |
| D  (Th_H) | MADS-box | <0.05 |
|  | GRF | <0.1 |
|  | ABI3VP1 |  |
